# Advanced Fabrication Protocol of an Elastic Porous Membrane for Organ-on-a-chip Applications

**DOI:** 10.64898/2026.02.26.708274

**Authors:** Nam Than, Hyun Jung Kim

## Abstract

Elastic porous membranes are essential components of mechanically active organ-on-a-chip and microphysiological system (MPS) platforms, where cyclic strain is required to recapitulate physiologically relevant tissue mechanics. However, existing fabrication methods are often difficult to reproduce, low throughput, or dependent on specialized infrastructure, limiting their adoption across laboratories. Many protocols also lack quality control steps for ensuring device assembling and reproducibility. In this paper, we present a robust and accessible fabrication and quality control workflow for the consistent production of elastic porous PDMS membranes. The method uses commercially available heat presses, release liners, and pre-patterned membrane wafers to enable rapid membrane molding. We describe a quality control framework, including visual verification of porous regions and wettability testing for surface activation, to ensure irreversible PDMS bonding and reliable device assembly. Together, this workflow improves fabrication yield, reduces device failure, and supports reproducible implementation of elastic porous membrane in organ-on-a-chip applications.

## INTRODUCTION

Traditional animal models often fail to recapitulate key aspects of human physiology, resulting in limited translational relevance and high attrition rates in clinical trials^1^. In addition, conventional *in vivo* testing is slow, costly, and ethically challenging. These limitations have motivated the development of advanced human-relevant *in vitro* microphysiological systems (MPS) for drug discovery^2^. While human-derived two-dimensional (2D) cell cultures and organoids offer biological relevance, they lack the dynamic mechanical and biochemical cues necessary to mimic native tissue function and host-microbiome interactions^3^. Organ-on-a-chip technology addresses these gaps by integrating human cells within microengineered systems that recreate the architecture, mechanics, and physiology of real organs. These MPS hold the potential to accelerate drug discovery, reduce development costs, and provide more predictive data for personalized medicine^3^.

The majority of MPS models require a basement membrane as a structural component to form and recapitulate a functional cellular unit on a biocompatible, flexible, and porous membrane. This membrane serves as a physical substrate for extracellular matrix (ECM) deposition, supporting the adhesion of anchorage-dependent cells while providing directional cues for cell polarization. Moreover, the membrane establishes a defined boundary between adjacent tissue compartments, permitting the co-culture of multiple cell types, such as epithelium and endothelium, under distinct microenvironments, including differential oxygenation. Membrane porosity enables molecular transport, supporting physiologically relevant processes including absorption^4, 5^ and efflux^6^, as well as cell transmigration across epithelial or endothelial barriers^7–9^. Membrane elasticity also allows it to function as a biomechanical interface, transmitting dynamic strain that mimics organ-specific motions such as intestinal peristalsis^10, 11^ or lung breathing^12^. The ability to model these structural, biochemical, and mechanical features positions the porous membrane as a key regulator of tissue organization and function in MPS platforms. Therefore, rapid and reproducible fabrication of high-quality porous PDMS membranes is essential to advancing organ-on-a-chip devices for next-generation biomedical research and drug development.

Various materials are used for porous membrane fabrication, including polycarbonate (PC)^13, 14^, polyethylene terephthalate (PET)^15^, polytetrafluoroethylene (PTFE)^16^, poly(lactic-co-glycolic acid) (PLGA)^17^, and polydimethylsiloxane (PDMS)^7, 10, 18–20^. Among these, PDMS is most widely utilized owing to its biocompatibility, optical transparency for real-time in situ observation, gas permeability, and elasticity that enables cyclic strain^21^. Cyclic strain is a critical feature of organ-on-a-chip systems because rhythmic mechanical deformation regulates cellular and immunological responses that are absent under static culture conditions. In a heart-on-a-chip model, cyclic strain induces antifibrotic signaling in cardiac fibroblasts^22^, while in a coronary artery-on-a-chip model, it elicits anti-inflammatory responses in the vascular endothelium^23^. In a lung alveolus-on-a-chip model, breathing-like motions suppress influenza virus replication and activate protective innate immune signaling through mechanosensitive pathways that are absent under static culture conditions^12^. Moreover, cyclic stretching facilitates greater particle movement within laminar flow regimes^11^ and may promote the clearance of excessive bacterial accumulation, supporting long-term host-microbe interaction studies^8^.

Several alternative elastic membrane materials have also been explored, including ECM hydrogel-based, synthetic ECM-based, or thermoplastic elastomer-based membranes. However, these approaches present limitations when mechanical actuation and long-term stability are required. Collagen- or fibrin-based membranes are mechanically fragile, prone to cracking when dried or swelling and creep when hydrated^24^, while also exhibiting bonding instability with microfluidic channel materials^25, 26^. Synthetic ECM membranes, such as gelatin methacrylate (GelMA) hydrogels, offer improved tunability^27^ but remain constrained due to limited stretchability^28^, variability in pore size, swelling ratio, and degradation rate^29^, as well as low fatigue resistance under cyclic stretching^30^. Thermoplastic polyurethane-based membranes have also been explored^31, 32^, but their broader adoption in organ-on-a-chip research has not yet been widely reported. In contrast, porous PDMS membranes offer high elasticity, fatigue resistance under cyclic deformation, and robust irreversible bonding to PDMS microfluidic channels, and are supported by mature, well-established fabrication workflows widely implemented in organ-on-a-chip platforms.

Despite its advantages, existing PDMS membrane fabrication methods face technical and practical limitations that restrict their scalability and broader adoption. For example, etching-based approaches for producing through-hole membranes typically require a chemical hood, multi-step processing involving chemical etching or chemical vapor deposition, and extensive hands-on technician effort^33^. Methods based on toluene-soluble polystyrene are also time-consuming and inevitably produce non-uniform pore structures and arrangements^34^, introducing confounding variability into downstream biological experiments. Additionally, fabricated membranes may exhibit a suboptimal bonding success rate during organ-on-chip device assembly^33^, further reducing effective yield and overall throughput. Compression-based approaches offer both conceptual and practical simplicity. However, current methods either operate at low throughput^19, 20^ or rely on patented pneumatic compression devices, which may restrict accessibility across laboratories^35^. Lastly, the lack of quality control (QC) disclosure in current fabrication protocols represents a critical omission, as key parameters such as pore openness and surface activation are rarely validated. Although tracer permeability assays can confirm molecular transport across the membrane, these assessments typically require a fully assembled device, making failures costly and time-intensive to identify. Implementing QC metrics before bonding would enable early detection of defective membranes and significantly improve fabrication yield, reproducibility, and throughput.

Overall, the field lacks rapid, scalable, and standardized fabrication methods that yield reproducible, quality-assured porous PDMS membranes. Addressing these challenges is critical to improving the reliability, throughput, and accessibility of organ-on-a-chip devices for drug discovery and translational biomedical research. Therefore, we present a new method for the rapid and scalable production of thin (≤10 μm thick) porous PDMS membranes and the associated QC process. Briefly, we have developed and validated a reproducible protocol for fabricating high-quality porous PDMS membranes using a commercial heat press, release liners, and a pre-patterned membrane wafer (Fig. 1A). Next, we describe a quality control procedure that enables routine verification of pore openness to ensure consistent membrane functionality (Fig. 1B). We also present considerations for release liner selection. Lastly, we demonstrate how the choice of release liner and surface activation method influences PDMS surface tension and bonding reliability between organ-on-a-Chip compartments (Fig. 1C).

**Fig. 1.**
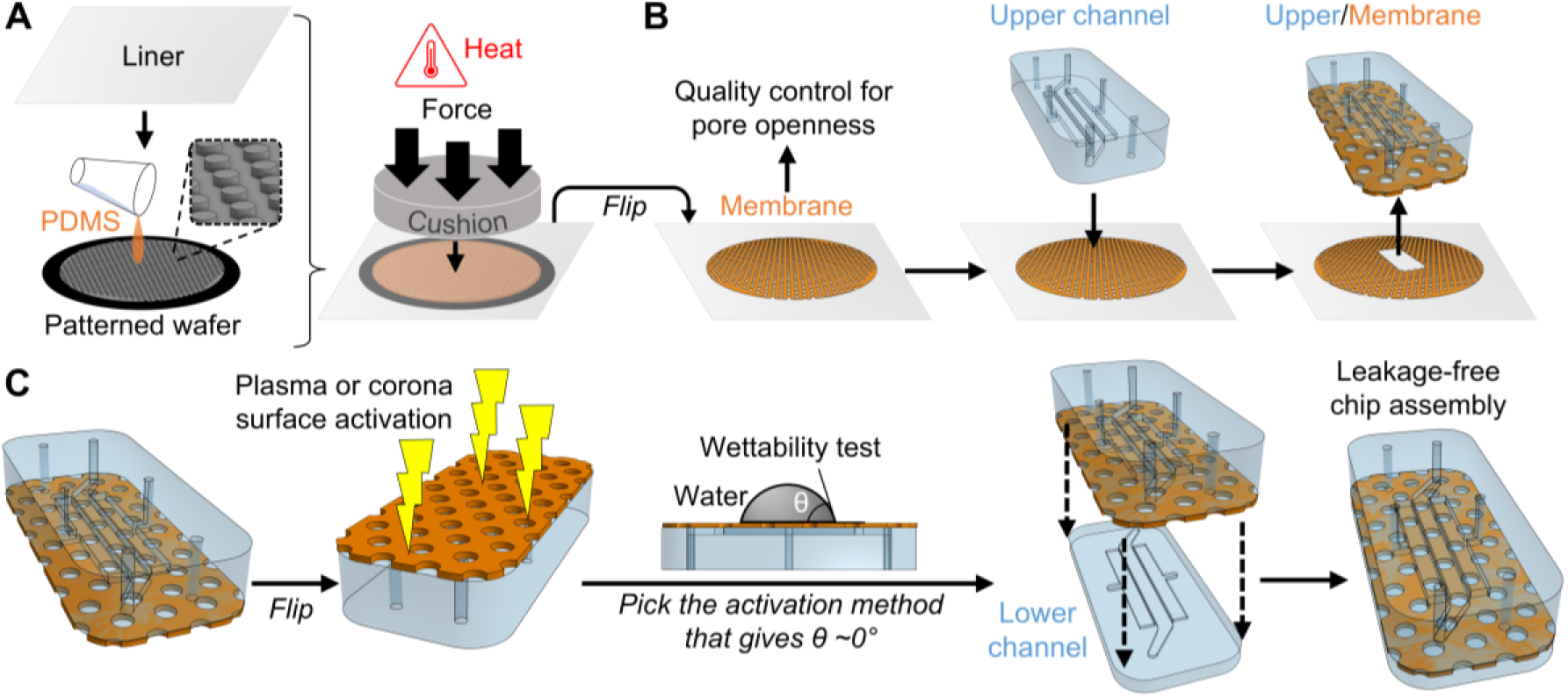
Schematic overview of the fabrication and quality control workflow for porous PDMS membranes. A. Major Step 1: The membrane is fabricated using a heat press that applies simultaneous compression and heat. A release liner facilitates both the transfer of the cured membrane from the silicon wafer and its subsequent integration with the upper channel compartment. B. Major Step 2: Quality control is conducted on the produced membrane to assess pore openness before bonding to the upper channel. Porous membrane illustrations are not to scale. C. Major Step 3: Quality control before fabrication to match the release liner with the appropriate surface-activation method, ensuring irreversible bonding during Organ-on-a-Chip assembly.

## MATERIALS

### Chemicals & materials

#### Polydimethylsiloxane (PDMS)

PDMS (Sylgard 184, Dow Corning) with a 10:1 (w/w) mixing ratio (elastomer base to curing agent) was used throughout this protocol.

#### Silicon membrane wafer

Each pre-patterned membrane wafer contains an array of micropillars with 10 μm height, 10 μm diameter, and 25 μm center-to-center spacing, arranged in an equilateral triangular lattice (Supplementary Fig. S1A). This pillar geometry corresponds to a membrane porosity of approximately 14.5%, calculated as the ratio of porous area within the triangular unit to the total area of that unit (Supplementary Fig. S1B). Membrane wafers can be fabricated using standard photolithography procedures in a clean room or purchased from a commercial vendor. Pillar patterns can be custom-designed with different geometries or porosities depending on experimental needs.

#### Transparent release liner

3M Scotchpak fluoropolymer-coated polypropylene release liner (9741 or 9744, 3M Technologies) was used. Alternative release-liner candidates include polyester (FRA344, Fox River Associates), polycarbonate sheet (85585K72, McMaster-Carr), or ethylene tetrafluoroethylene (ETFE) sheet (2539N18, McMaster-Carr). These alternatives typically require additional surface pre-activation before use, as described in the Troubleshooting section.

#### Ethanol

200-proof ethanol (C961Y21, Thomas Scientific) was used.

### Equipment

#### Heat press

A clamshell heat press (HPN-CRAFT-1515, HeatPress Nation) was used. The heat press features a heating element in the upper platen and a central pressure-control knob, both essential for uniform heating and compression.

#### Optical microscope

An inverted light microscope (DMi1, Leica) equipped with a 10× objective, numerical aperture (NA) 0.22 (506271, Leica), and a 20× objective, NA 0.33 (506272, Leica), was used for routine inspection and quality control of membrane pore openness. A stereoscope (MDG41, Leica) equipped with 1× or 1.6× objectives and bright- or dark-field illumination was used during the alignment and bonding of the upper and lower gut chip layers.

#### Plasma cleaner

A plasma cleaner (COVANCE-1MPR, Femto Science Inc.) was used for rapid surface activation of PDMS and for irreversible bonding between PDMS components.

#### Corona discharge treater

A handheld corona discharge treater (080-1203-1, Electro-Technic Products) was used when the selected release liner inhibits plasma-induced surface activation (see Major Step 3), providing an alternative activation method for reliable PDMS bonding.

#### Contact angle goniometer

The following items can be used for constructing a low-cost, do-it-yourself (DIY) contact angle goniometer: a white-light table lamp (1920K113, McMaster-Carr), a polycarbonate sheet (85585K33, McMaster-Carr), and a smartphone or low-cost digital camera (see Major Step 3).

#### Vacuum desiccator

A vacuum desiccator (F42025-0000, Bel-Art) was used for degassing the PDMS mixture.

#### Additional tools

These tools were useful to have: a seam roller (62395T13, McMaster-Carr), a rigid plastic card (38EV78, Grainger), a ¼-inch borosilicate glass plate (8476K23, McMaster-Carr), aluminum foil (01-213-105, FisherBrand), and an abrasion-resistant polyurethane rubber sheet with ultra-smooth texture (2185T31, McMaster-Carr).

### Software

Contact angle measurement can be performed using ImageJ (NIH; https://imagej.net/ij/), a free, open-source image-processing program. The “Angle Tool” can be used to quantify the droplet boundary from images captured using the DIY goniometer.

## PROCEDURE

### Major Step 1: Heat press-assisted rapid fabrication of PDMS porous membrane

#### Timing: 2 hours

Our protocol utilizes a commercially available clamshell heat press capable of providing conformal compression and heating. Conceptually, PDMS will be cured between a pre-patterned silicon wafer and a release liner under compression and heat (Fig 2A). Pores form only when the release liner is in conformal contact with the tops of the patterned pillars. To achieve uniform compression, an elastic cushion (3 mm thick) such as a slab of polyurethane (PU) is placed above the release liner (Fig. 2B). PU resists adhesion to excess PDMS and is easily cleaned. Alternatively, a silanized PDMS slab can achieve the same performance.

**Fig. 2.**
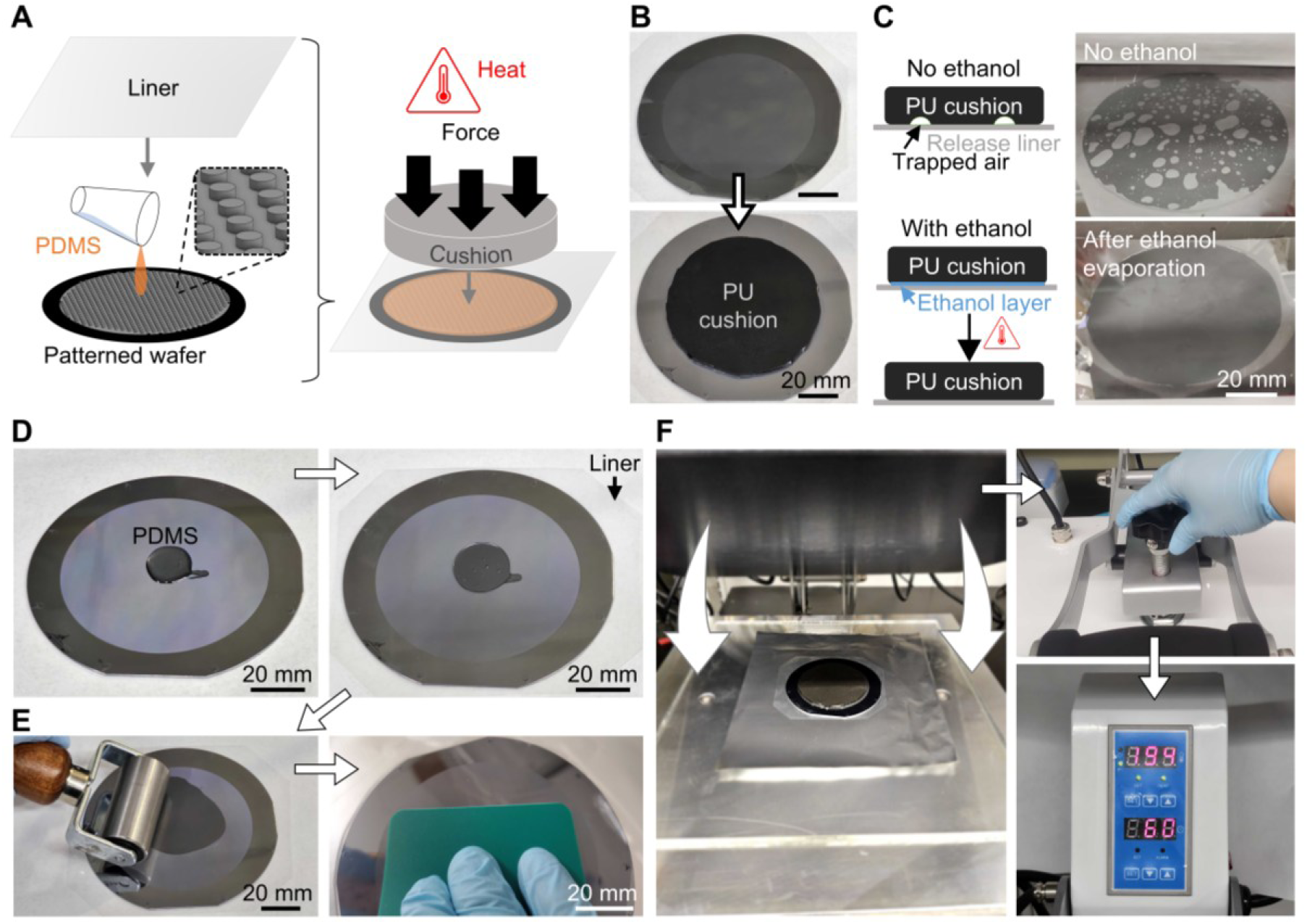
Heat press-assisted rapid fabrication of PDMS porous membrane. A. Schematic illustration of the fabrication principle, showing the combined roles of heat, compression, elastic cushion, and a pre-patterned silicon wafer. B. Placement of a polyurethane cushion prior to compression and heating. C. Application of ethanol layer to eliminate trapped air between the liner and cushion. D. Deposition of PDMS and placement of the release liner onto the membrane wafer. E. Spreading of PDMS using a seam roller and a plastic card to achieve uniform film thickness. F. Compression and baking process.

It is also essential to prevent air entrapment between the cushion and the release liner by wetting the release liner with 100% ethanol before cushion placement (Fig. 2C, top schematic and top-right image). Trapped air can lead to non-uniform compression. Ethanol’s low surface tension allows it to wet the cushion efficiently and displace trapped air. Upon ethanol evaporation during heating, the two surfaces remain in conformal contact (Fig. 2C, bottom schematic and bottom-right image). No interference with PDMS curing was observed under the tested conditions. This major step establishes the uniform compression and controlled thermal curing necessary for reproducible formation of open pores across the PDMS membrane.

1. Prepare PDMS by combining the elastomer base and curing agent at the desired ratio. Mix well and degas in a desiccator for 1 hour before use.
2. Prepare the platen by placing a borosilicate glass sheet on the lower platen to provide a flat, abrasion-resistant surface. Cover the glass with a wrinkle-free sheet of aluminum foil for easy cleanup.
3. Place the membrane wafer in the center of the platen.
4. Dispense PDMS by pouring 1 g of PDMS onto the center of the wafer to initially cover approximately 5-10% of the surface (Fig. 2D, left panel).
5. Apply the release liner by placing a pre-cut transparent release liner onto the PDMS (Fig. 2D, right panel).

a. Pre-activation of the liner surface may be required (see Troubleshooting).
6. Use a seam roller and a rigid plastic card to smoothly and evenly spread PDMS across the patterned area (Fig. 2E). Excess PDMS should be forced outward.
7. Discontinue scraping once the membrane is evenly coated and no additional PDMS is expelled.
8. Wet the top surface of the liner with a uniform layer of 100% ethanol, either by spraying or by pipetting approximately 500 µL of ethanol and evenly distributing it across the surface using a rigid plastic card.
9. Place an elastic polyurethane cushion above the release liner, with the ultra-smooth surface facing the liner (Fig. 2B).

a. Optional: Pre-cut the cushion to match the patterned area.
10. Release the pressure knob before lowering the top platen to avoid accidental crushing of the wafer.
11. Apply compression by lowering the platen and tightening the pressure knob until firm resistance is reached (Fig. 2F).
12. Cure the membrane by turning on the heating. Temperature and duration may be optimized based on the thermal stability of the selected materials.

a. For a PU cushion: 90 °C for 60 minutes.
b. For a PDMS cushion: 120 °C for 40 minutes.
13. Turn off the heat when the set time is reached, release the pressure knob, and slowly raise the upper platen.
14. Proceed to the membrane quality control step immediately.

##### Critical

Use mixed PDMS base/curing agent within 2 hours.

##### Critical

Do not use excessive downward force during Step 6 as the membrane wafer may break.

##### Critical

Do not use excess ethanol during Step 8 to avoid mixing with uncured PDMS.

##### Critical

Avoid non-knob clamshell heatpresses (lever-only systems) due to severe compression non-uniformity.

### Major Step 2: Quality control for membrane pore openness

#### Timing: 10 minutes

Once fabrication is complete, the porous PDMS membrane is detached from the silicon wafer by gently peeling off the release liner. The cured PDMS membrane preferentially adheres to the release liner rather than the wafer, enabling clean, damage-free separation. The membrane, still attached to the release liner, can then undergo the QC step to assess pore openness and overall structural integrity.

Quality control is a critical component of any manufacturing process. While SEM imaging can serve as proof of successful porous membrane fabrication^33^, SEM is expensive and time-consuming, rendering it impractical for routine use during regular production. We introduce a practical and rapid alternative for assessing pore openness using a standard inverted light microscope. The evaluation is based on a clear physical distinction: open pores produce a jagged, serrated tear edge when the membrane is gently torn, whereas unopened pores, resulting from incomplete conformal contact between the release liner and micropillars, yield a straight tear line (Fig. 3A). This method, termed the serration check, can be performed at random points along a pre-marked cushion boundary to confirm local pore openness (Fig. 3B). However, this approach is destructive and provides only spot-based verification, necessitating another QC step for assessing pore openness across the entire membrane area.

**Fig. 3.**
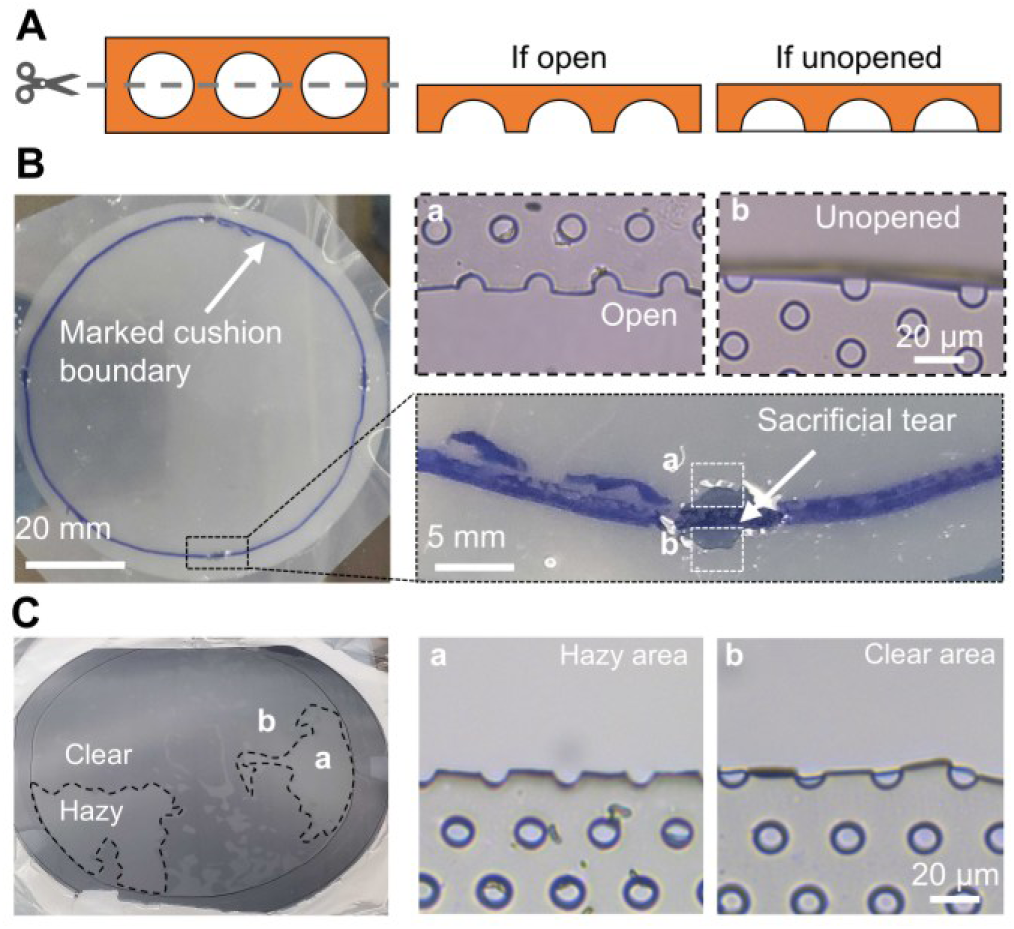
Quality control for membrane openness. A. Schematic illustration of the serration-check principle, showing that open pores produce a serrated edge upon tearing, whereas unopened pores yield a straight edge. B. Serration check performed by optically inspecting a sacrificial tear at the marked cushion boundary, demonstrating distinct membrane morphologies with and without compression. The region beneath the cushion (panel a) exhibits a serrated edge indicative of open pores, whereas the region outside the cushion (panel b) shows a smooth, straight edge consistent with a non-porous membrane. C. An intentionally low-yield example demonstrates that open-pore regions exhibit a hazy or matte appearance, whereas unopened regions remain transparent and reflect the gray color of the wafer. A serration check performed in each region verifies the association between porosity and hazy appearance.

Through repeated observations, we noted that immediately after fabrication and removal of the cushion, and before peeling the membrane-bound release liner, open-pore regions appear visually hazier than unopened regions when viewed on the wafer (Fig. 3C; also see Anticipated Results). Thus, this major step integrates global visual assessment with targeted serration checks to provide a robust, high-confidence QC workflow for confirming membrane pore openness.

15. Use a marker to trace the boundary of the cushion. Gently lift off the cushion without disturbing the liner (Fig. 3B).
16. Inspect the liner and mark the boundary of the hazy/matte region after a 2-min cooling period (Fig. 3C). These regions likely correspond to open-pore areas.
17. Peel off the liner gently, ensuring that the porous PDMS membrane remains adhered to the liner rather than the wafer.

a. Optional: Apply a small amount of 100% ethanol between the PDMS and wafer to aid detachment
18. Remove residual PDMS from the wafer. The wafer can be reused once cooled to room temperature.

a. Optional: Use 100% ethanol for more thorough cleaning, followed by drying overnight in a 60 °C oven.
19. Use tweezers to create small sacrificial scratches/tears at random points along both the cushion boundary (Step 15) or the hazy/matte boundary (Step 16).
20. Examine these sacrificial tears under a 10× and 20× objective. Open-pore regions display clear serrated edges, whereas unopened regions show straight, smooth tears (Fig. 3B, zoom-in panels).
21. Refine the marked boundaries based on the serration check results.
22. Dry and store the membrane at 60 °C until ready for organ-on-a-chip assembly.

##### Critical

If the membrane remains stuck to the wafer rather than the liner, refer to the Troubleshooting section.

##### Critical

The hazy/matte contrast is most reliable immediately after a short cooling period (Step 16); partial detachment of the membrane-bound liner, extended delay, or observation while the membrane remains hot may reduce visibility.

##### Note

Patchy, hazy regions indicate insufficient or non-uniform compression; refer to the Limitations section.

### Major Step 3: Quality control for surface activation of the PDMS membrane

#### Timing: 2 h

Following QC of membrane pore openness, the membrane is activated and bonded to the upper channel of the Organ Chip device using standard methods^10, 19^. However, we discovered that PDMS surfaces cured in contact with fluoropolymer-coated release liners resist plasma activation and instead require corona discharge for complete activation (Fig. 4A & 4B, liner-facing side). This plasma-activation inhibition effect was observed only in the fluoropolymer-coated liners, whereas other liners can be successfully activated by both methods (Fig. 4C). Failure to properly activate the PDMS surface compromises bonding integrity and can lead to liquid leakage during operation of organ-on-a-chip devices. Thus, we propose a QC step to verify successful surface activation on both the wafer-facing and the liner-facing side of the membrane. This QC step is accomplished by a simple wettability test and contact angle measurement.

23. Prepare a test sample by cutting a 1 cm × 1 cm piece of the porous membrane.
24. Activate the wafer-facing side of the sample using a plasma cleaner (125 W, 75 s).
25. Apply a 20-µL droplet of deionized water to the activated surface and capture an image using a goniometer (DIY option provided below).

a. Successful activation is indicated by near-complete spreading of the droplet, producing a contact angle of 0°-25° (empirically determined), whereas a higher angle indicates incomplete or failed activation.
b. If activation is successful, proceed to Step 24.
c. If activation fails, repeat Steps 23-25 with a corona discharge treater (2 minutes) instead of a plasma cleaner.
26. Activate both the Organ Chip upper channel part and the porous membrane using the validated activation method.
27. Align and bond the upper part to the membrane. Incubate at 60°C overnight or 80°C for 1 h.
28. Trace the perimeter of the upper part with a razor blade to separate it from the surrounding membrane, then peel the membrane-bound upper part off the release liner.

a. Refer to the Troubleshooting section if the membrane does not transfer cleanly from the liner to the upper channel part.
29. Perform the same surface activation QC (Steps 23-25) on the liner-facing side of the upper-channel-bound membrane (Fig. 4A-B).
30. Activate both the upper-channel-bound membrane and the bottom channel using the validated activation method.
31. Align and bond the two compartments under a stereoscope. Incubate at 60°C overnight or 80°C for 1 h.
32. Confirm irreversible bonding by gently flexing and pressing the device edges. Properly bonded interfaces will remain fully sealed without detachment.
33. Flush DI water through both channels to confirm a leak-free bond.

**Fig. 4.**
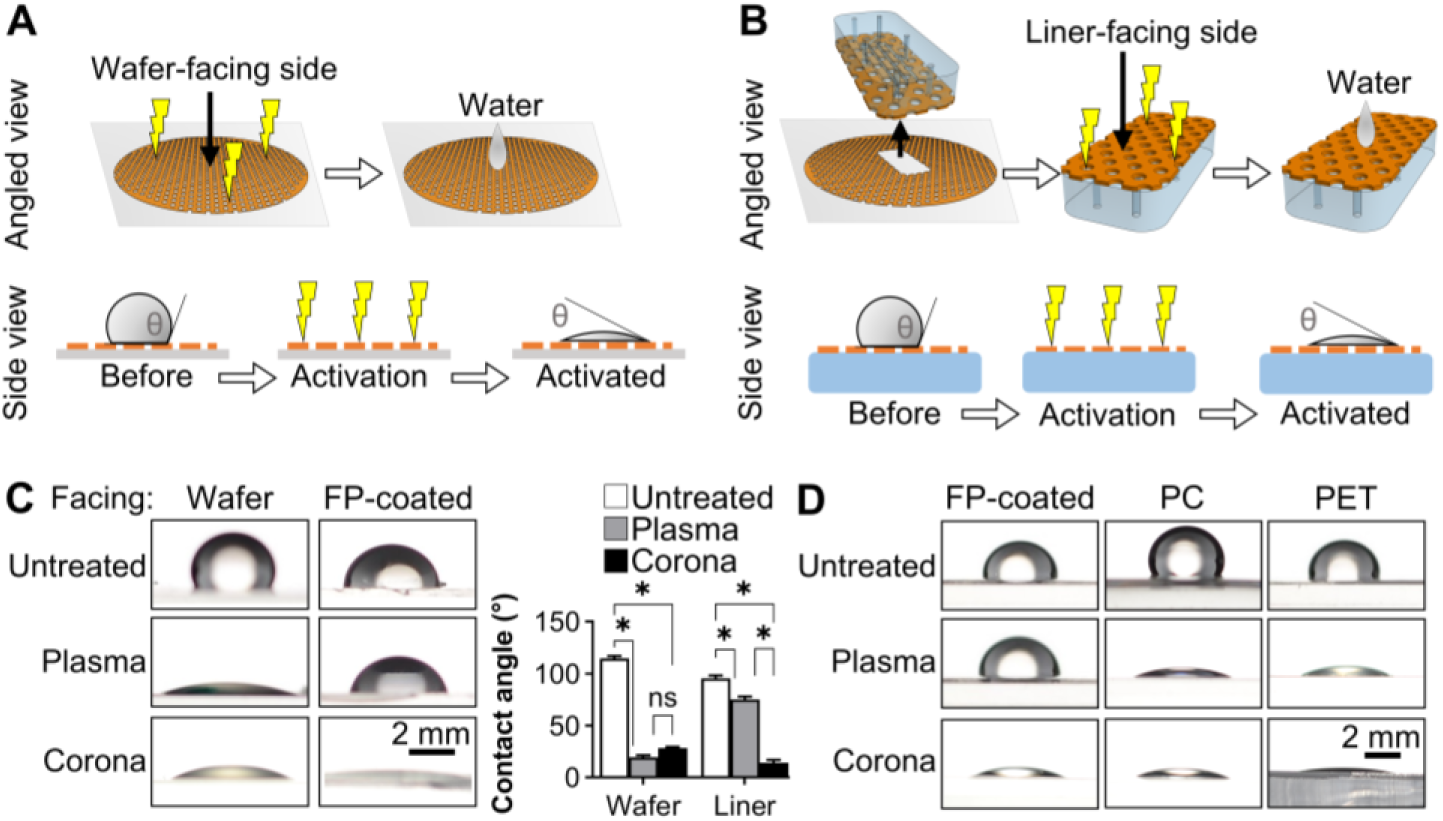
Quality control for surface activation of the PDMS membrane. A. Schematic illustration of the wettability test for evaluating surface activation on the wafer-facing side of the porous membrane. The angled view depicts the experimental procedure, while the side view illustrates the expected wettability outcome. A low contact angle indicates successful activation. B. Schematic illustration of the wettability test for evaluating surface activation on the liner-facing side of the porous membrane. The angled view depicts the experimental procedure, while the side view illustrates the expected wettability outcome. A low contact angle indicates successful activation. C. The wafer-facing side of the membrane can be activated by either plasma or corona treatment, whereas the fluoropolymer (FP) liner-facing side requires corona discharge exclusively. \**p* < 0.001; ns, not significant. D. Representative contact-angle measurements demonstrating that plasma-activation inhibition is specific to the fluoropolymer-coated liner series.

##### Critical

This major step should be performed for every new release liner to properly match the activation method with the specific release liner, thus safeguarding downstream irreversible, leakage-free Organ Chip assembly.

##### Optional

We developed a DIY goniometer constructed from readily available laboratory materials because a commercial contact angle goniometer can be costly (Supplementary Fig. S2A). A strong white-light source (such as a table lamp) is positioned behind a diffuser to generate uniform, parallel illumination. A polycarbonate sheet can be used as a simple diffuser. The sample is placed in front of the diffuser, aligned to the same vertical height as the camera’s focal plane. A smartphone or low-cost digital camera is mounted at a fixed distance equal to its focal distance. Images are acquired with the droplet edge in sharp focus using the camera’s automatic exposure settings. If a phone camera is used, modest digital zoom (e.g., 2×) can improve edge resolution. Under this back-illumination, a water droplet appears with a sharp, dark perimeter at a high resolution, enabling accurate contact-angle measurement using freely available image-processing software such as ImageJ (Supplementary Fig. S2B).

## TROUBLESHOOTING

The release liner serves two critical roles: first, it sandwiches the PDMS layer between the membrane wafer and itself during fabrication for pore formation, and second, it facilitates detachment of the cured PDMS membrane from the silicon wafer (i.e., primary transfer). Subsequently, during a secondary transfer, the PDMS membrane must be completely released from the liner and bonded to the desired PDMS chip component (e.g., the upper channel layer). A successful release liner must therefore demonstrate balanced adhesion properties. It should adhere more strongly to the PDMS membrane than to the silicon wafer, ensuring efficient primary transfer, while remaining sufficiently non-adhesive to allow clean detachment during the secondary transfer.

This delicate balance in adhesion strength requires extensive empirical optimization. Through systematic testing, we have identified several release liners, namely the 3M Scotchpak fluoropolymer-coated release liners 9741 and 9744, that exhibit optimal adhesion profiles for both primary and secondary transfers without requiring surface pretreatment. However, if the user employs a different type of release liner, the following problems could arise.

### Problem 1: Primary transfer fails

The membrane remains on the silicon wafer and does not transfer to the release liner after peeling.

#### Potential solution

Plasma activation of the release liner surface can increase adhesion when working with high-release materials such as polycarbonate or ETFE, thereby improving primary transfer yield. However, plasma activation effectiveness is time- and humidity-sensitive, which can introduce variability associated with surface hydrophobic recovery. To mitigate these effects, ambient humidity should be monitored and maintained within a consistent range, and bonding of PDMS components must be performed immediately following plasma treatment, thereby ensuring reproducibility across fabrication batches.

### Problem 2: Secondary transfer fails

The membrane transfers successfully to the liner (primary transfer) but then remains adhered to the liner and fails to bond to the PDMS upper channel layer.

#### Potential solution

Case A: Primary transfer required surface activation (from Problem 1)

Excessive plasma activation can cause permanent bonding between the liner and PDMS. Activation parameters (e.g., duration, power) should be fine-tuned to achieve just enough adhesion for primary transfer while still allowing membrane release during secondary transfer.

Case B: The release liner has intrinsically high adhesion

If the liner binds strongly even without pre-activation, apply surface passivation, such as silanization with Trichloro(1H,1H,2H,2H-perfluorooctyl)silane (448931, Sigma-Alrich), to increase release force and enhance clean membrane detachment.

## DATA ANALYSIS

Contact-angle measurements were analyzed using ImageJ. Briefly, the Angle Tool was used by first drawing a baseline parallel to the substrate at the base of the water droplet, followed by a second line tangent to the droplet’s edge (Supplementary Fig. S2B). Each droplet yielded two contact angles (*θ*′ and *θ*″), corresponding to the left and right interfaces. The final contact angle (*θ*) was calculated as the average of *θ*′ and *θ*″.

## ANTICIPATED RESULTS

Major Step 1 is expected to consistently and rapidly produce high-yield porous membrane areas, which are then verified in Major Step 2. Serration checks should clearly differentiate between open and unopened pores. For example, as demonstrated in Fig. 3B, pores directly under the cushion present a serrated tear edge (panel a), confirming efficient pore opening, whereas pores outside the compressed region exhibit a straight tear edge (panel b).

We further confirmed that the hazy/matte regions observed immediately after fabrication correspond to open-pore membrane areas, validated through serration checks. A similar visual cue was briefly noted but not discussed in another reported protocol^35^. Even though the precise mechanism was not investigated, this verified guideline ensures that only the porous membrane region is used in the downstream assembling steps.

Having identified the porous membrane region, Major Step 3 is expected to address any release-liner interferences with surface activation. In our protocol, fluoropolymer-coated liners consistently reduce plasma activation efficacy but remain responsive to corona treatment. Although the precise mechanism was not investigated, and this observation may not generalize to all fluoropolymer-coated release liner products, it underscores the importance of matching release liner surface chemistry with the appropriate surface activation method to ensure robust bonding and leakage-free Organ Chip assembly.

## LIMITATIONS

Compared to existing reported protocols, our pipeline is advantageous in one or more of these aspects: (a) higher porous area yield per unit of time per wafer, (b) better uniformity, or (c) better bonding efficiency. However, several technical limitations must be addressed for further scaling up the manufacturing process. First, although commercial heat presses can accommodate platens as large as 20 × 20 in, such dimensions are not necessarily compatible with silicon membrane wafers of the same size. Silicon wafers are inherently brittle, and handling oversized substrates increases the risk of fracture during alignment or compression.

Second, a uniform pressure distribution across the platen is also a limiting factor. Heat presses equipped with a single central pressure knob may produce non-uniform compression, with pressure decreasing toward the platen edges. In our experiments, 90-100% open-pore yield was consistently achieved using 3-in-diameter round wafers (or, 41.0-45.6 cm^2^/h/wafer), while 3 × 7 in rectangular wafers achieved approximately 80-90% (108.4-121.9 cm^2^/h/wafer). Alternatively, commercial hydraulic heat presses equipped with dual heating platens and four corner guide rods to maintain platen parallelism could potentially offer improved compression uniformity and are expected to restore high pore-opening yield across larger wafers.

Lastly, manual corona treater use introduces operator-dependent variability and is less efficient for batch processing. In contrast, plasma treatment systems support high-throughput workflows, enabling simultaneous activation of multiple samples with minimal labor. For scaling up membrane manufacturing, a release liner that does not inhibit plasma activation is preferred, as it allows consistent and automated surface activation across multiple chips. Looking forward, an ideal release liner should satisfy two criteria: (a) balanced adhesion strength sufficient for both primary and secondary membrane transfer (as detailed in the Troubleshooting section), and (b) compatibility with plasma activation to support scalable, reproducible production. If both criteria cannot be met simultaneously, balanced adhesion strength should take priority over activation convenience, as reliable membrane transfer is a fundamental requirement.

## ACKNOWLEDGMENTS

This work was supported in part by the Cleveland Clinic Foundation Catalyst SPARK award, the Kenneth Rainin Foundation Innovator Awards, the Crohn’s and Colitis Foundation of America Senior Research Awards, the Cleveland Clinic VeloSano Pilot Grants, and the NIH NCI IMAT program (1R33CA286797).

## AUTHOR CONTRIBUTIONS

N.T. and H.J.K. conceptualized the study, performed the experiments, analyzed the data, and wrote the manuscript.

## DECLARATION OF COMPETING INTERESTS

The authors declare no competing interests.

## Notes

### Competing Interest Statement

The authors have declared no competing interest.

## References

1. Sun, D.; Gao, W.; Hu, H.;, et al. Why 90% of clinical drug development fails and how to improve it? Acta Pharmaceutica Sinica B 2022, 12, 3049–3062.

2. Ingber, D. E. Human organs-on-chips for disease modelling, drug development and personalized medicine. Nat. Rev. Genet. 2022, 23, 467–491.

3. Shin, Y. C.; Than, N.; Min, S.;, et al. Modelling host-microbiome interactions in organ-on-a-chip platforms. Nat. Rev. Bioeng. 2024, 2, 175–191.

4. Maschmeyer, I.; Lorenz, A. K.; Schimek, K.;, et al. A four-organ-chip for interconnected long-term co-culture of human intestine, liver, skin and kidney equivalents. Lab Chip 2015, 15, 2688–2699.

5. Kim, H. J.; Ingber, D. E. Gut-on-a-Chip microenvironment induces human intestinal cells to undergo villus differentiation. Integr Biol 2013, 5, 1130–40.

6. Blundell, C.; Yi, Y. S.; Ma, L.;, et al. Placental drug transport-on-a-chip: a microengineered in vitro model of transporter-mediated drug efflux in the human placental barrier. Adv Healthc Mater 2018, 7, 1700786.

7. Huh, D.; Matthews, B. D.; Mammoto, A.;, et al. Reconstituting organ-level lung functions on a chip. Science 2010, 328, 1662–1668.

8. Kim, H. J.; Li, H.; Collins, J. J.;, et al. Contributions of microbiome and mechanical deformation to intestinal bacterial overgrowth and inflammation in a human gut-on-a-chip. Proc Natl Acad Sci U S A 2016, 113, E7–15.

9. Shin, W.; Kim, H. J. Intestinal barrier dysfunction orchestrates the onset of inflammatory host-microbiome cross-talk in a human gut inflammation-on-a-chip. Proc Natl Acad Sci U S A 2018, 115, E10539–E10547.

10. Kim, H. J.; Huh, D.; Hamilton, G.;, et al. Human gut-on-a-chip inhabited by microbial flora that experiences intestinal peristalsis-like motions and flow. Lab Chip 2012, 12, 2165–2174.

11. Shin, Y. C.; Shin, W.; Koh, D.;, et al. Three-Dimensional Regeneration of Patient-Derived Intestinal Organoid Epithelium in a Physiodynamic Mucosal Interface-on-a-Chip. Micromachines (Basel) 2020, 11, 663.

12. Bai, H.; Si, L.; Jiang, A.;, et al. Mechanical control of innate immune responses against viral infection revealed in a human lung alveolus chip. Nat. Commun. 2022, 13, 1928.

13. Wu, J.; Zhang, B.; Liu, X.;, et al. An Intelligent Intestine-on-a-Chip for Rapid Screening of Probiotics with Relief-Enteritis Function. Adv. Mater. 2024, 36, 2408485.

14. Shah, P.; Fritz, J. V.; Glaab, E.;, et al. A microfluidics-based in vitro model of the gastrointestinal human–microbe interface. Nat. Commun. 2016, 7, 11535.

15. Kulthong, K.; Duivenvoorde, L.; Sun, H.;, et al. Microfluidic chip for culturing intestinal epithelial cell layers: Characterization and comparison of drug transport between dynamic and static models. Toxicology in Vitro 2020, 65, 104815.

16. Sellgren, K. L.; Hawkins, B. T.; Grego, S. An optically transparent membrane supports shear stress studies in a three-dimensional microfluidic neurovascular unit model. Biomicrofluidics 2015, 9.

17. Na, N.; Kim, M.; Kim, J.;, et al. Fabrication and Customization of Highly Porous PLGA Membranes Utilizing Near-Field Electrospinning, Thermal Transitions, and Multilayer Strategies. Adv. Eng. Mater. 2024, 26, 2400917.

18. Low, L. A.; Mummery, C.; Berridge, B. R.;, et al. Organs-on-chips: into the next decade. Nature Reviews Drug Discovery 2021, 20, 345–361.

19. Shin, W.; Kim, H. J. 3D in vitro morphogenesis of human intestinal epithelium in a gut-on-a-chip or a hybrid chip with a cell culture insert. Nat. Protoc. 2022, 17, 910–939.

20. Kefallinou, D.; Grigoriou, M.; Boumpas, D. T.;, et al. Fabrication of a 3D microfluidic cell culture device for bone marrow-on-a-chip. Micro and Nano Engineering 2020, 9, 100075.

21. Leung, C. M.; De Haan, P.; Ronaldson-Bouchard, K.;, et al. A guide to the organ-on-a-chip. Nature Reviews Methods Primers 2022, 2, 33.

22. Gartner, T. C. B.; Wang, Y.; Leiteris, L.;, et al. Cyclic strain has antifibrotic effects on the human cardiac fibroblast transcriptome in a human cardiac fibrosis-on-a-chip platform. J. Mech. Behav. Biomed. Mater. 2023, 144, 105980.

23. Hou, Y.; Ziakas, G.; Hopkins, T.;, et al. Human coronary artery tri-culture organ-chip recapitulates anti-inflammatory effect of pulsatile wall strain. bioRxiv 2025, 2025.09. 26.678578.

24. Meyer, M. Processing of collagen based biomaterials and the resulting materials properties. Biomed. Eng. Online 2019, 18, 24.

25. Lei, F.; Liang, M.; Liu, Y.;, et al. Multi-compartment organ-on-a-chip based on electrospun nanofiber membrane as in vitro jaundice disease model. Advanced Fiber Materials 2021, 3, 383–393.

26. Dabaghi, M.; Shahriari, S.; Saraei, N.;, et al. Surface modification of PDMS-based microfluidic devices with collagen using polydopamine as a spacer to enhance primary human bronchial epithelial cell adhesion. Micromachines 2021, 12, 132.

27. Shirahama, H.; Lee, B. H.; Tan, L. P.;, et al. Precise tuning of facile one-pot gelatin methacryloyl (GelMA) synthesis. Sci. Rep. 2016, 6, 31036.

28. Montazerian, H.; Baidya, A.; Haghniaz, R.;, et al. Stretchable and bioadhesive gelatin methacryloyl-based hydrogels enabled by in situ dopamine polymerization. ACS applied materials & interfaces 2021, 13, 40290–40301.

29. Aljaber, M. B.; Verisqa, F.; Keskin-Erdogan, Z.;, et al. Influence of gelatin source and bloom number on gelatin methacryloyl hydrogels mechanical and biological properties for muscle regeneration. Biomolecules 2023, 13, 811.

30. Zhu, J.; Wei, Y.; Yang, G.;, et al. Peptide-based rigid nanorod-reinforced gelatin methacryloyl hydrogels for osteochondral regeneration and additive manufacturing. Nat. Commun. 2025, 16, 7090.

31. Mitta, E.; Gilmore, A.; Malliri, A.;, et al. A stretchable and biomimetic polyurethane membrane for lung alveolar in vitro modelling. Sci. Rep. 2025, 15, 14585.

32. Domansky, K.; Leslie, D. C.; McKinney, J.;, et al. Clear castable polyurethane elastomer for fabrication of microfluidic devices. Lab Chip 2013, 13, 3956–3964.

33. Quirós-Solano, W.; Gaio, N.; Stassen, O.;, et al. Microfabricated tuneable and transferable porous PDMS membranes for Organs-on-Chips. Sci. Rep. 2018, 8, 13524.

34. Yap, J. H.; Zhang, H.; Okamura, Y.;, et al. Thin microporous polydimethylsiloxane membrane prepared by phase separation and its applications for cell culture. Materialia 2024, 38, 102247.

35. Novak, R.; Didier, M.; Calamari, E.;, et al. Scalable Fabrication of Stretchable, Dual Channel, Microfluidic Organ Chips. J. Vis. Exp. 2018, e58151.

