## Supplemental Information 1 for "Advanced Fabrication Protocol of an Elastic Porous Membrane for Organ-on-a-chip Applications"

**Nam Than[1] and Hyun Jung Kim[1,2,*]**
[1]Department of Inflammation and Immunity, Cleveland Clinic, Cleveland, OH 44195, USA
[2]Cleveland Clinic Lerner College of Medicine of Case Western Reserve University, Cleveland, OH 44195, USA

***Correspondence to**:
Hyun Jung Kim, PhD
Department of Inflammation and Immunity
Cleveland Clinic Research, Cleveland Clinic
9500 Euclid Ave., NE3
Cleveland, OH 44195, USA


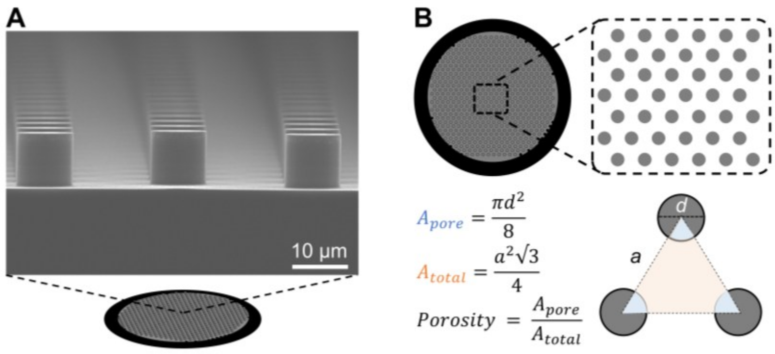


**Fig. S1. Design and porosity calculation of a membrane wafer**

A. Scanning electron micrograph (SEM) of the micropillar array on the silicon wafer used for pore formation. The micrograph was provided by the manufacturer.

B. Schematic top-view of the pillar array and the calculation of membrane porosity based on pillar geometry and spacing. In this design, the total area of the equilateral triangle colored in orange is calculated as $\frac{a^2\sqrt{3}}{4}$. The porous area within this triangle is the sum of the 3 sectors colored in blue and calculated as $\frac{\pi d^2}{8}$. Porosity is the ratio of the porous area over the total area. a, center-to-center distance between any 2 pillars; d, diameter of the pillar.

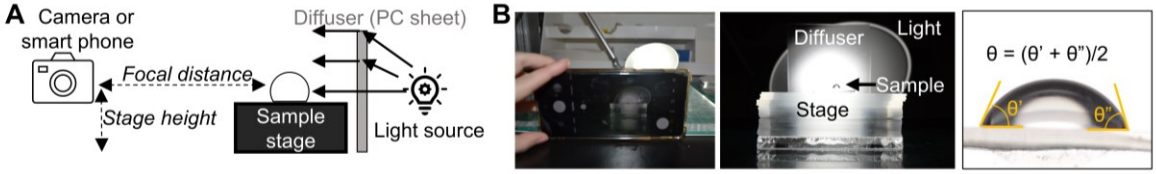


**Fig. S2. A low-cost, do-it-yourself contact angle goniometer set-up**

A. Schematic of the DIY contact-angle goniometer for surface wettability measurements, assembled from readily available laboratory components, including a table lamp, a polycarbonate diffuser, and a smartphone or digital camera with an auto-exposure setting. The camera must be positioned at a focal distance that produces a sharp focus of the droplet edge.

B. Representative high-resolution droplet image acquired using the DIY goniometer and corresponding contact-angle analysis. For each droplet, two contact angles ($\theta'$ and $\theta''$) were measured at the left and right solid–liquid–air interfaces, and the reported contact angle ($\theta$) was calculated as the average of $\theta'$ and $\theta''$.
